# Investigating the role of APOE: limitations of human pluripotent stem cell-derived cerebral organoids

**DOI:** 10.1101/2020.11.30.405654

**Authors:** Damián Hernández, Louise A. Rooney, Maciej Daniszewski, Lerna Gulluyan, Helena H. Liang, Anthony L. Cook, Alex W. Hewitt, Alice Pébay

## Abstract

Apolipoprotein E (APOE) is the most important susceptibility gene for late onset of Alzheimer’s disease, with the presence of APOE-ε4 associated with increased risk of developing Alzheimer’s disease. Here, we reprogrammed human fibroblasts from individuals with different *APOE-ε* genotypes into induced pluripotent stem cells, and generated isogenic lines with different APOE profiles. We then differentiated these into cerebral organoids for six months and assessed the suitability of this *in vitro* system to measure APOE, β amyloid, and Tau phosphorylation levels. We identified intra- and inter-variabilities in the organoids’ cell composition. Using the CRISPR-edited *APOE* isogenic lines, we observed more homogenous cerebral organoids, and similar levels of APOE, β amyloid, and Tau between the isogenic lines, with the exception of one site of Tau phosphorylation which was higher in the APOE-ε4/ε4 organoids. These data describe that pathological hallmarks of AD are observed in cerebral organoids, and that their variation is mainly independent of the *APOE-ε* status of the cells, but associated with the high variability of cerebral organoid differentiation. It demonstrates that the batch-to-batch and cell-line-to-cell-line variabilities need to be considered when using cerebral organoids.

## Introduction

Alzheimer’s disease (AD) is a chronic and progressive disease that leads to the degeneration of neurons key to the function of the brain and to cognitive decline. Histological hallmarks of AD consist in extracellular dense amyloid plaques composed of β amyloid (Aβ); and intracellular neurofibrillary tangles of hyper-phosphorylated Tau protein (for review see [1, 2]). Plaques and tangles hamper neuronal and synaptic functions leading to cell death, associated with cognitive decline [2]. Despite enormous research efforts, there are still no definitive treatments for AD. Studying AD is hindered by the lack of non-invasive means for obtaining brain tissue from living donors. This can now be overcome by using human induced pluripotent stem cells (iPSCs). *In vitro* cultures recapitulate fundamental events observed in AD, and iPSC-based studies have confirmed findings obtained in other models [3–7]. Further, the development of human cerebral organoids is bringing a novel level of organisation to *in vitro* models. These organoids are considered as recapitulating the first months of human neural development, with presence of neural stem/ progenitor cells, neurons and astrocytes as well as appearance of structured layers and regionalisation [8–10]. Cerebral organoids also possess features of mature neurons, such as dendritic spines and electrical activity [11]. Hence, those technological advances are starting to address the pressing need for accurate and complex human disease models to improve our molecular understanding of neurodegenerative diseases including AD [12, 13].

Apolipoprotein E (APOE) is a protein involved in the regulation of cholesterol metabolism [14]. In the central nervous system, APOE is mainly produced and secreted by astrocytes [15]. There are three APOE isoforms in humans, APOE2, APOE3 and APOE4, which differ from one another in residues 112 and 158, and respectively encoded by alleles *APOE-ε2* (Cys112, Cys158), *APOE-ε3* (Cys112, Arg158) *or APOE-ε4* (Arg112, Arg158). It is now unequivocally established that *APOE* is the most important susceptibility gene for late onset of AD, with the presence of *APOE-ε4* associated with increased risk of developing AD [16–19]. Using human iPSC-derived cells, various reports have shown association of the *APOE4* variant with impaired cellular phenotypes [7]. In neural progenitors, *APOE-ε4/ε4* is associated with an earlier neuronal differentiation [20, 21]. In astrocytes, the APOE4 variant is associated with cholesterol accumulation and impaired Aβ uptake [21, 22]. In microglial-like cells, APOE4 is linked to a reduced Aβ phagocytosis [21], an increased cytokine release, and impaired metabolism, phagocytosis and migration [23]. In neurons, *APOE-ε3/ε4* is linked to increased cell death and increased phosphorylated Tau [24] while *APOE-ε4/ε4* is associated with an increased number of synapses [21], increased GABAergic neuron degeneration [22], increased production of Aβ [21, 22] and increased amount of phosphorylated Tau [22]. Similar phenotypes have been observed in six-month-old cerebral organoids, absent at earlier time points, and explained by the absence of APOE expression in younger organoids [21]. It was also recently reported that cerebral organoids derived from AD patients with an *APOE4/4* genotype recapitulate aspects of AD *in vitro*, in particular increased production of Aβ and amount of phosphorylated Tau, with these effects absent In *APOE4* cerebral organoids derived from non AD patients [25]. Here, we generated iPSCs from individuals with different *APOE* risk genotypes (*APOE-ε3/ε3* and *APOE-ε4/ε4*) as well as a CRISPR-edited isogenic line of *APOE-ε4/ε4* to *APOE-ε3/ε3*, differentiated those to cerebral organoids and monitored the key pathological events of AD (Aβ deposits and Tau phosphorylation) after six months. Altogether our data suggests that heterogeneity of cerebral organoids remains a major issue for *in vitro* modelling of specific phenotypes observed in AD.

## Materials and Methods

### Ethics

All experimental work performed in this study was approved by the Human Research Ethics committees of the Royal Victorian Eye and Ear Hospital (11/1031H), University of Melbourne (1545394), University of Tasmania (H0014124) with the requirements of the National Health & Medical Research Council of Australia (NHMRC) and conformed with the Declaration of Helsinki [26].

### iPSC generation and maintenance

iPSCs were generated using skin fibroblasts obtained from subjects over the age of 18 years with no diagnosis of dementia, by episomal method as described [27] (Table 1). Briefly, reprogramming was performed on passage 2 fibroblasts by nucleofection (Lonza Amaxa Nucleofector) with episomal vectors expressing *OCT4, SOX2, KLF4, L-MYC, LIN28* and shRNA against *p53* [28] in feeder- and serum-free conditions using TeSR-E7 medium (Stem Cell Technologies). Subsequently, reprogrammed colonies were manually dissected to establish clonal cell lines [27] for the iPSC lines MBE2960 [29], MBE2968 (clone 1 [23] and clone 2 in this paper), TOB0002 (clone 3 [23] and clones 1 and 5 in this paper), whilst the lines TOB0064 and TOB0121 were reprogrammed and selected as a polyclonal line using an automated platform as described in [30]. All lines were subsequently expanded and characterised. The iPSCs were maintained manually in serum-free and feeder-free conditions using StemFlex and vitronectin (ThermoFisher), as described in [31]. Medium was changed every second day and cells were passaged weekly when colonies reached 80% confluency.

**Table 1.** iPSC lines used.

| **ID** | **Age at biopsy** | **Gender** | ***APOE* status** | **Clones** | **Reference** |
| --- | --- | --- | --- | --- | --- |
| **TOB0002** | 52 | M | *APOE4/4* | c1 | This paper (Figure 1) |
|  |  |  |  | c3 | [23] |
|  |  |  |  | c5 | This paper (Figure 1) |
| **TOB0064** | 73 | M | *APOE4/4* | polyclonal | This paper (Figure 2) |
| **TOB0121** | 68 | F | *APOE4/4* | polyclonal | This paper (Figure S1) |
| **MBE2960** | 78 | M | *APOE3/3* | c1 | [29] |
|  |  |  |  | c2 | [29] |
|  |  |  |  | c3 | [29] |
| **MBE2968** | 65 | F | *APOE3/3* | c1 | [23] |
|  |  |  |  | c2 | This paper (Figure S2) |
| **iTOB0064** | 73 | M | *APOE3/3* | 2 | This paper (Figure 2) |
|  |  |  |  | 3 |  |

### Generation of isogenic lines

Genome editing of the TOB0064 iPSC line was performed with the CRISPR/Cas9 system. Single guide (sg) RNA sequence was designed as described by Zhang’s laboratory [32]. Briefly, 20 bp sgRNA with PAM sequence NGG closer to the DNA targeting region of interest were designed using the CRISPR tool from the website Benchling and selected based on the highest on-target and off-target score [32, 33]. For homology directed repair, a single strand (ss) DNA of 100 bp was designed with the point mutation of interest in the middle of the sequence. The *APOE-ε4* allele of the TOB064 line was genetically modified to generate isogenic lines with a homozygous *APOE-ε3* allele (APOE3/3 isogenic lines) with the following sequence of sgRNA and the ssDNA: sgRNA (5’- /AltR1/rGrCrG rGrArC rArUrG rGrArG rGrArC rGrUrG rCrGrG rUrUrUrUrArG rArGrC rUrArU rGrCrU /AltR2/ -3’), ssDNA (5’- TGCGCAAGCTGCGTAAGCGGCTCCTCCGCGATGCCGATGACCTGCAGAAG**t**GCCTGGCAGTGTACCAGGCCGGGGCCCGCGAGGGCGCCGAGCGCGGCCT-3’). The ribonucleoprotein (RNP) complex consisting of Cas9 protein and sgRNA (containing a tracrRNA labelled with a red fluorophore ATTO 550) was assembled in a duplex buffer (all from Integrated DNA Technologies). RNP complex was subsequently transfected into 100,000 dissociated iPSCs, resuspended in a final volume of 13 μL in buffer R (Invitrogen) by electroporation (1200 V, 30 ms, 1 pulse) with the Neon transfection system (Invitrogen). After electroporation the cells were immediately plated at low density onto vitronectin-plated 6 well plates containing warm Stemflex media supplemented with ROCK inhibitor (RevitaCell, Gibco). After 48 hours, ATTO 550 positive cells were sorted by flow cytometry (BD) and plated at low density onto vitronectin-plated 6-well plates. Individual clones were manually picked and screened for SNP editing by PCR and Sanger sequencing (Australian Genome Research Facility). One colony was subsequently dissociated, genotyped as to obtain a pure edited population. Two clones were then amplified and used in the study (Table 1). The resultant isogenic lines were named iTOB0064_2_ and iTOB0064_3_.

### Virtual karyotype

Copy number variation (CNV) analysis of original fibroblasts and iPSCs was performed using Illumina HumanCore Beadchip arrays as we described [31]. CNV analyses were performed using PennCNV and QuantiSNP [34, 35] with default parameter settings. Chromosomal aberrations were deemed to involve at least 10 contiguous single nucleotide polymorphisms (SNPs) or a genomic region spanning at least 1MB [34, 35]. The B allele frequency (BAF) and the log R ratio (LRR) were extracted from GenomeStudio (Illumina, USA) for representation.

### Differentiation to the three germ layer

Embryoid bodies (EB) were obtained as described [36] and using tri-lineage differentiation kit (Stem Cell Technologies). Germ layer differentiation was assessed by immunochemistry.

### Cerebral organoid differentiation

Differentiation into cerebral organoids was performed with a commercially available differentiation kit from Stemcell Technologies (Catalog number 08570-1) based on previously reported protocols [8, 9]. Briefly, iPSCs were dissociated and 9,000 cells were seeded into each well of a U-shape 96-well plates in EB Formation Medium (Basal Medium 1 with Cerebral Organoid Supplement A) and 50 mM Y27632 (Tocris). The medium was changed every second day without Y27632. After 5 days in culture, organoids were transferred into an ultra-low attachment 24-well plate containing Induction Medium (Basal Medium 1 with Cerebral Organoid Supplement B). On day 7, organoids were embedded into a droplet of Matrigel and furthered cultured in Expansion Medium (Basal Medium 2 with Cerebral Organoid Supplements C and D) in ultra-low attachment 6-well plates. At day 10, medium was changed to Maturation Medium (Basal Medium 2 with Cerebral Organoid Supplement E) and plates were placed onto an orbital shaker within the incubator. The shaking speed was 65 rpm (relative centrifugal force of 0.11808*g*). Medium was changed every 2-3 days. Organoids were cultured and harvested at 6 months (180 days).

### Immunocytochemistry

Immunocytochemistry was performed using the following primary antibodies: mouse anti-OCT3/4 (2.5 μg/ml, sc-5279, Santa Cruz Biotechnology), mouse anti-TRA-1-60 (1 μg/ml, ab16288, Abcam), mouse anti-NESTIN (10 μg/ml, AB22035, Abcam), rabbit anti-alpha-fetoprotein (AFP, 10 μg/ml, st1673, Sigma-Aldrich), and mouse anti-smooth muscle actin (SMA, 10 μg/ml, MAB1420, R&D Systems). Cells were then immunostained with isotype-specific secondary antibodies (Alexa Fluor 568 or 488, Life Technologies). Nuclei were counterstained using DAPI (Sigma-Aldrich) and mounted in Vectashield (Vector Labs). Specificity of the staining was verified by the absence of staining in negative controls consisting of the appropriate negative control immunoglobulin fraction (Dako). Images were acquired on a Zeiss AxioImager M2 fluorescent microscope using ZEN software (Zeiss).

### Western blotting

Cerebral organoids were lysed in RIPA buffer (Thermo Fisher Scientific) containing a cocktail of protein and phosphatases inhibitors (Thermo Fisher Scientific). Proteins were denatured in NuPAGE sample reducing buffer and boiled for 3 minutes and heated up to 85°C for 2 minutes. 50 μg of proteins was separated by SDS-PAGE using 4–12% Novex Tris-Glycine gels in Novex Tris-Glycine SDS running buffer (all from ThermoFisher). Proteins were then transferred onto a polyvinylidene difluoride membrane (ThermoFisher), which was blocked with SuperBlock (Thermo Fisher Scientific; 1 hour at room temperature or overnight at 4°C), washed in phosphate-buffered saline containing 0.1% Tween 20 (PBS-T) and followed by incubation with primary antibodies or negative isotype controls diluted in blocking buffer for 1 hour (Covance PRB-278P rabbit polyclonal IgG anti-PAX6, 10μg/mL; Dako Z0334 rabbit polyclonal IgG anti-GFAP, 2μg/mL; Abcam ab18207 rabbit polyclonal IgG anti-β-Tubulin III, 0.4μg/mL; Thermo Fisher Scientific 701241 rabbit monoclonal IgG anti-APOE, 0.5μg/mL, Abcam ab8224 mouse monoclonal IgG anti-β-Actin, 1μg/mL; Dako P0448 rabbit IgG isotype control, 1μg/mL and Dako P0260 rabbit monoclonal IgG anti-β-Actin, 10μg/mL; rabbit IgG isotype control, 1μg/mL) at room temperature. After a second wash in PBS-T, the membranes were incubated with rabbit anti-mouse (Invitrogen 10500C, 130ng/mL) or goat anti-rabbit (Invitrogen 10400C, 7.5ng/mL) secondary antibodies for 1 hour at room temperature. Before incubation with β-Actin antibody, each membrane was stripped from the previous antibody by incubation with Restore Western Blot Stripping Buffer (Thermo Fisher Scientific) for 15 to 30 min at room temperature followed by a PBS-T wash and blocking with SuperBlock for 30 min. The membranes were scanned with the ChemiDoc MP Imaging System (Biorad), and protein band intensities were determined by computerized densitometry (Image Lab 6.1, Biorad) and expressed as fold change relative to the control after normalization to β-actin. Expected molecular weights for PAX6: 46-48 kDa; β-Tubulin: 50-55 kDa, GFAP: 37-50 kDa, and β-Actin: 42 kDa.

### ELISA

Levels of total APOE protein were analysed by Human ELISA Kit from Invitrogen (Apolipoprotein E Human ELISA Kit). Levels of Aβ were analysed by ELISA (Human β Amyloid 42 ELISA Kit Wako high sensitive #298-64401 and Human β Amyloid 40 ELISA Kit Wako #298-64601) following the manufacturer’s instructions. Total Tau, phospho Tau (pS396, pS199, pT231 and pT181) levels were analysed by Human ELISA Kit from Invitrogen. Cerebral organoids and supernatants (for Aβ) were homogenized in RIPA buffer with phosphatase and protease inhibitors (Pierce, Thermo Fisher Scientific) collected and diluted in a dilution buffer provided by the manufacturer. All samples were normalised for protein concentration prior to the ELISA. A SPARK plate reader (TECAN) was used to quantify absorbance from the ELISA signals (OD 450 nm for all assays).

### Statistical analysis

All sets of experiments were performed at least three times in triplicates, unless specified (n refers to the number of independent experiments performed on different cell cultures). Data were expressed as mean ± standard error of the mean (SEM). Significance of the differences was evaluated using t-test (comparing two isogenic *APOE* lines) or one ANOVA followed by the Tukey’s test for multiple comparisons (multiple lines). Statistical significance was established at *p<0.05, **p<0.01, ***p<0.001, ****p<0.0001. All statistical analyses and graphical data were generated using Graphpad Prism software (v5.04, www.graphpad.com).

## Results

The cohort included iPSCs with specific *APOE* genotypes, generated from unrelated individuals: three *APOE-ε4/ε4* (APOE4/4: TOB0002, TOB0064, TOB0121), two *APOE-ε3/ε3* (APOE3/3: MBE2960, MBE2968) and one isogenic line *APOE3/3* (iTOB0064) developed by gene editing of the parental iPSC line TOB0064 (Table 1). Patients fibroblasts were reprogrammed to iPSCs by nucleofection of episomal vectors containing *OCT4, SOX2, KLF4, L-MYC, LIN28*, and shRNA against *p53*. The iPSC line MBE2960 was described in [29], MBE2968 clone 1 and TOB0002 clone 3 were described in [23]. Here, we report the characterisation of TOB0002 clones 1 and 5 (Figure 1), TOB0064 (Figure 2), TOB0121 (Figure S1), and MBE2968 clone 2 (Figure S2). These lines express markers of pluripotency, were able to differentiate to the three germ layers and were karyotypically normal (Figures 1, 2, S1, S2). The *APOE-ε* status was confirmed for each line by Sanger DNA sequencing (Figures 1, 2, S1, S2).

**Figure 1.**
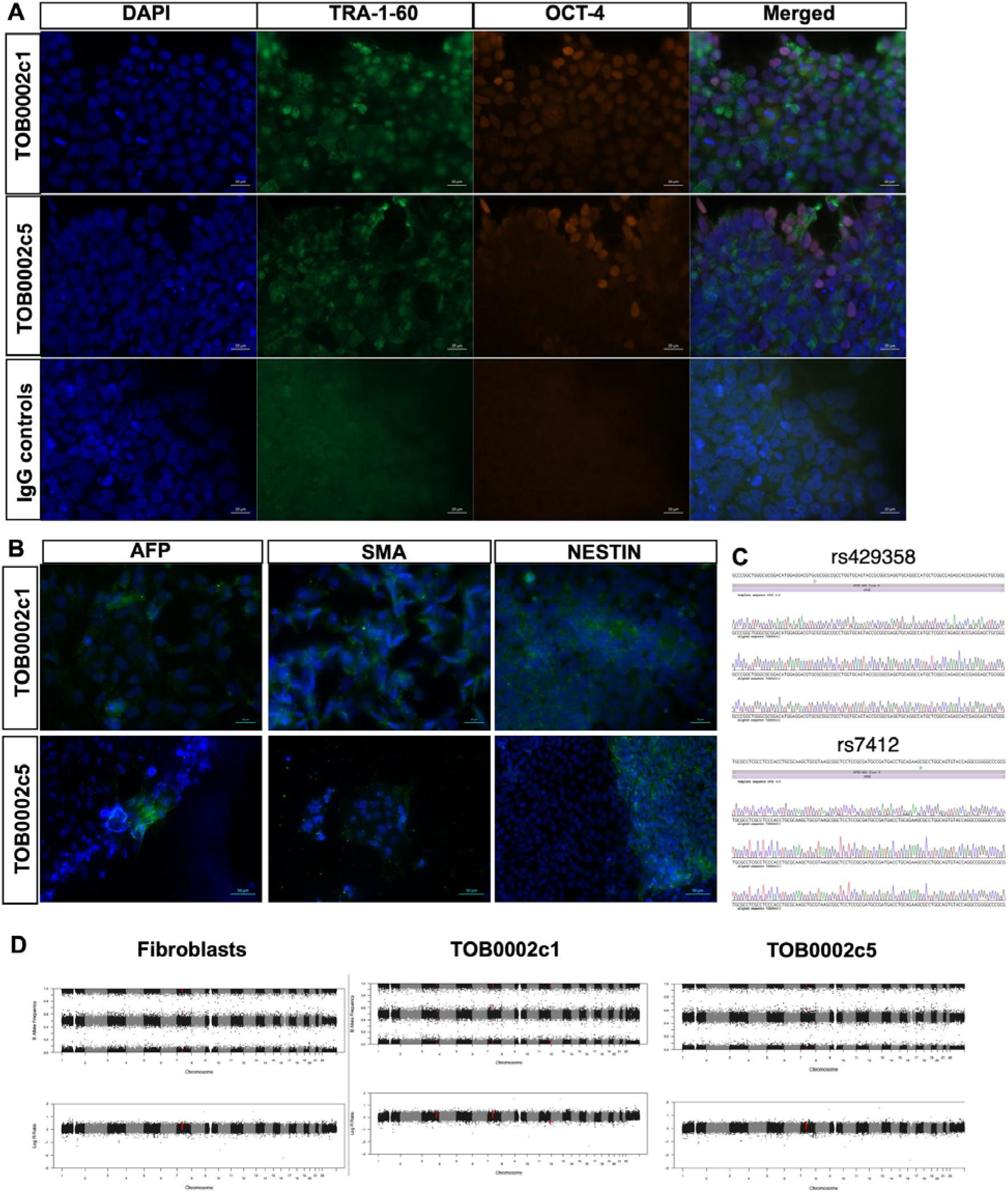
Characterisation of TOB0002. (**A**) **Generation of iPSC lines.** Representative images of TOB0002 clones (c) 1 and 5 showing expression of the pluripotency markers TRA-1-60 and OCT4, with DAPI counterstained and merged. Negative IgG controls are also displayed. (**B**) **Representative germ layer immunostaining of TOB0002.** TOB0002c1 and c5 demonstrate pluripotency by positive staining for markers of each embryonic germ layer; endoderm (AFP), mesoderm (SMA) and ectoderm (NESTIN). (**C**) ***APOE* genotyping.** Sanger sequencing of *rs429358* and *rs7412* in TOB0002c1, c3 and c5 confirms *APOE4/4* status. Note that c3 genotype was already published in [23] (**D**) **Copy Number Variation Analysis of TOB0002 in their original fibroblasts and iPSCs (p8).** Each panel shows the B allele frequency (BAF) and the log R ratio (LRR). BAF at values others then 0, 0.5 or 1 indicate an abnormal copy number. Similarly, the LRR represents a logged ratio of “observed probe intensity to expected intensity”. A deviation from zero corresponds to a change in copy number.

**Figure 2.**
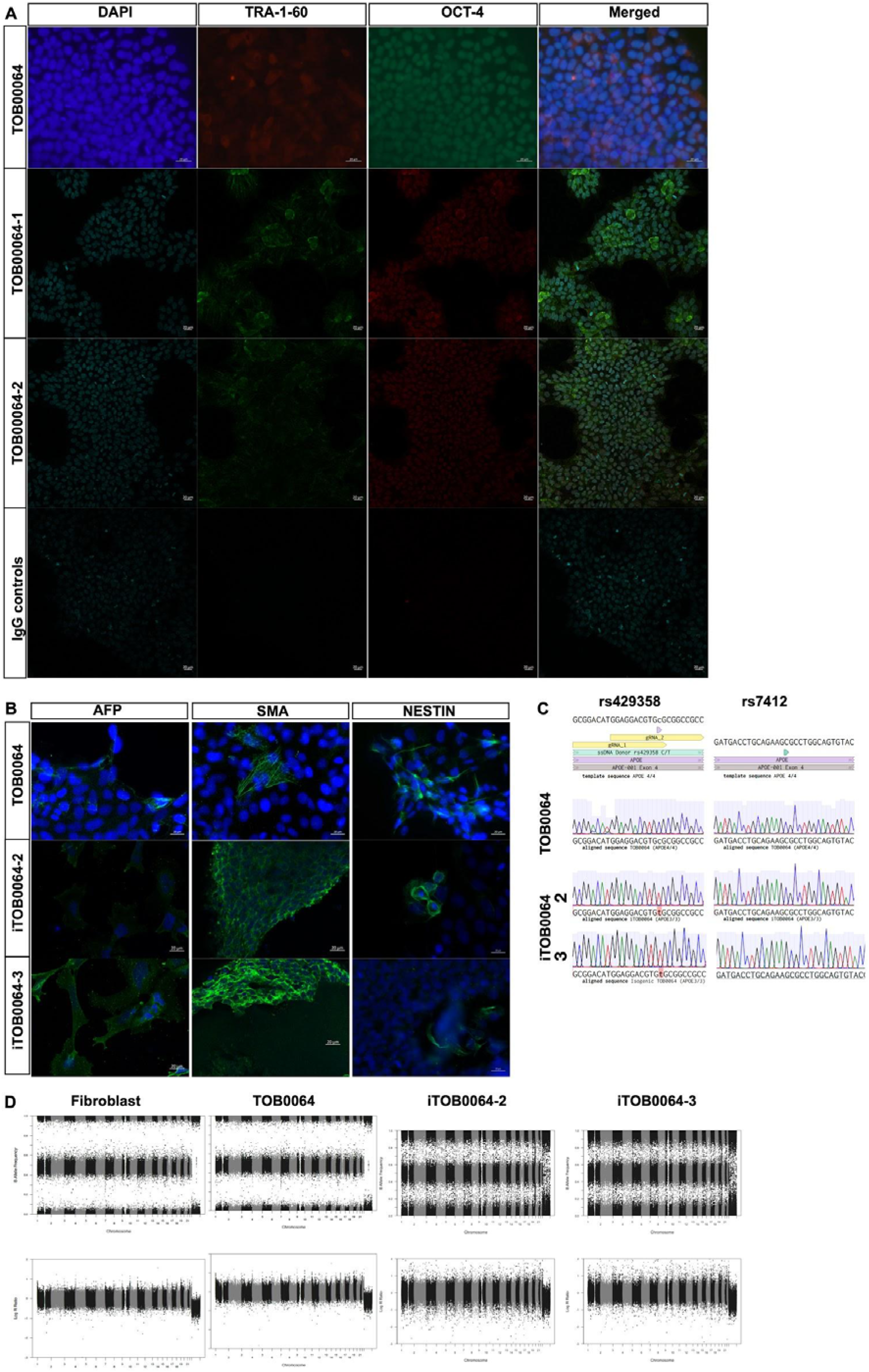
Characterisation of TOB0064 and iTOB0064. **(A) Generation of iPSC lines.** Representative images of TOB0064 showing expression of the pluripotency markers TRA-1-60 and OCT4, with DAPI counterstained and merged. **(B) Representative germ layer immunostaining of TOB0064.** Representative images of TOB0064 and iTOB0064_2,3_ differentiated to endoderm (AFP), mesoderm (SMA) and ectoderm (NESTIN). **(C)** *APOE* **genotyping.** Sanger sequencing of *rs429358* and *rs7412* confirms *APOE4/4* status in TOB0064 and APOE3/3 in iTOB0064_2,3_. Template sequence indicates *APOE4/4* with location of guide (g)RNAs **(D) Copy Number Variation Analysis of TOB0064 in their original fibroblasts, iPSCs and isogenic lines (iTOB0064_2,3_) (p8).** Each panel shows the B allele frequency (BAF) and the log R ratio (LRR). BAF at values other than 0, 0.5 or 1 indicate an abnormal copy number. Similarly, the LRR represents a logged ratio of “observed probe intensity to expected intensity”. A deviation from zero corresponds to a change in copy number.

All lines were differentiated to cerebral organoids for 6 months and quantification of cell type-specific proteins was assessed by western blots, using antibodies against PAX6 (neural progenitors), βIII-tubulin (neurons), and GFAP (astrocytes). For each iPSC line of either *APOE4/4* (TOB0002, TOB0064, TOB0121) or *APOE3/3* (MBE2960, MBE2968) genotypes, we first assessed the impact of differentiation batches on the levels of the cell type-specific proteins by analysing three independent batches of differentiation per cell line. All lines formed cerebral organoids in each batch of differentiation (Figure S3), with the exception of TOB0121 for which only one batch of differentiation was successful. In the western blots described in Figure 3A, each lane consisted of one batch of a cell line differentiation including 5 pooled organoids which was then quantified by densitometry analysis. Although each differentiation batch from all lines resulted in cerebral organoids, the western blot analysis indicated variations in levels of expression of PAX6, β-tubulin, and GFAP in 6 month-old organoids, mainly between cell lines (inter-cell line variability), yet also between batches of differentiation of each cell line (intra-cell line variability) (Figures 3A, S4). Given the observed heterogeneity between cell lines, this experimental setup could not be used to home in on a potential effect of APOE on cerebral organoid differentiation. To remove the inter-cell line variability and to investigate a potential impact of APOE on the cells’ differentiation potentials, we used the isogenic CRISPR-edited iPSC lines TOB0064 and iTOB0064 and differentiated these into 6 month-old cerebral organoids to quantify their cellular content by western blot (Figure 3B). For these experiments, TOB0064 was differentiated alongside iTOB0064. The TOB0064 differentiation batches were independent of those shown in Figure 3A. As prior, these lines were differentiated in three independent batches and the impact of each batch on cellular content was assessed. Variabilities in the expression of PAX6, β-tubulin and GFAP were observed across batches of differentiation but their levels were similar across the *APOE4/4* and *APOE3/3* organoids (Figures 3B, S5). This data thus indicates that the *APOE* genotype does not impact the ability of iPSCs to differentiate into neural cells and cerebral organoids. This lack of effect could however be masked if solely using unrelated iPSC lines instead of isogenic lines, as larger variations in cellular content were observed between unrelated cell lines with different APOE status, yet likely linked to other parameters than the *APOE* status. Altogether, this data confirms variability in the composition of cerebral organoids, as already reported by others [11, 37]. It describes both intra- and inter-cell line variations in organoid generation reproducibility per differentiation rounds. This data also demonstrates that the largest part of the variability in differentiation of iPSCs towards organoids is cell line-specific, and can thus be reduced by using isogenic lines.

**Figure 3.**
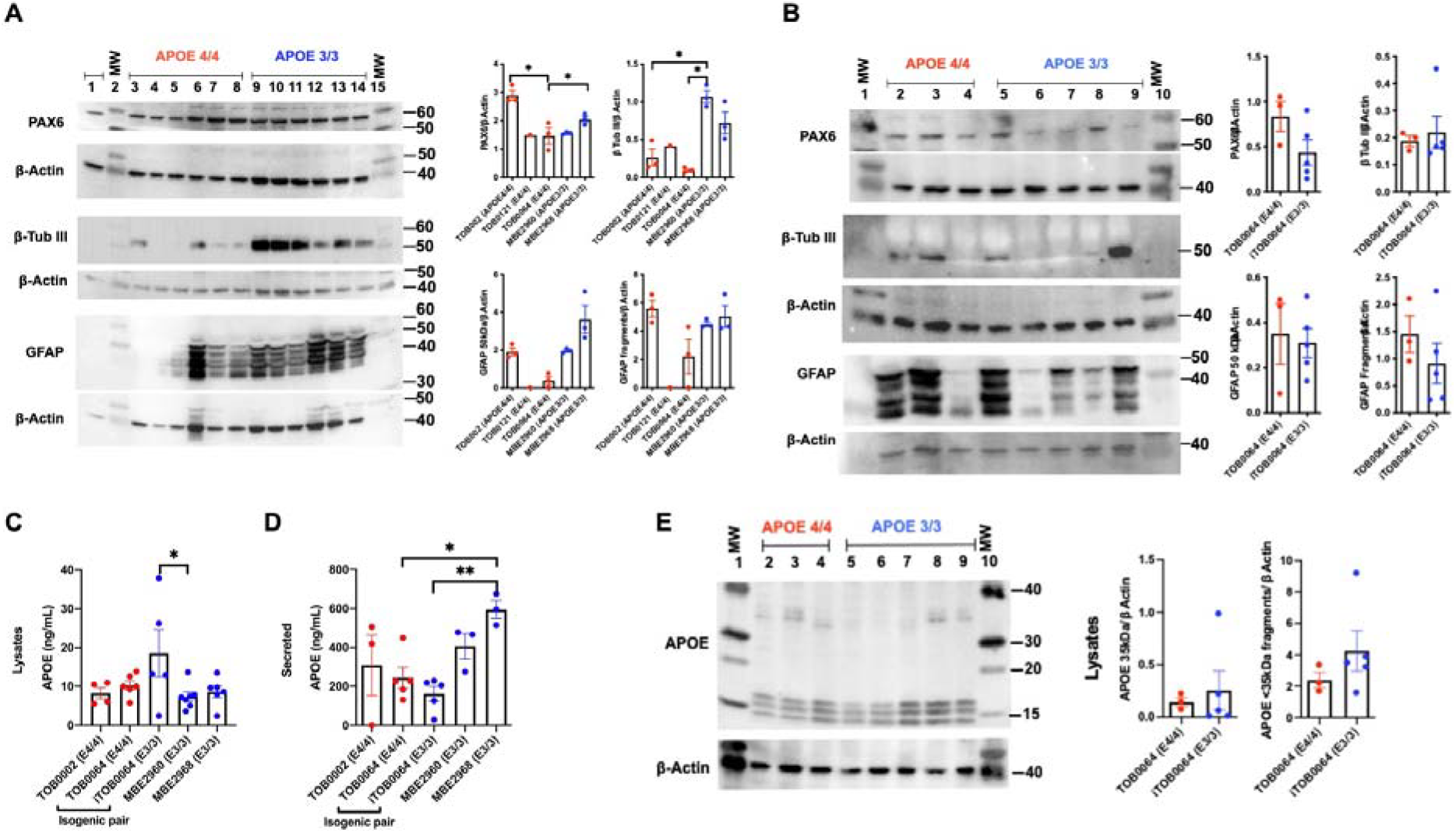
Heterogeneity of cerebral organoids. (**A, B**) Western blots of PAX6, β-tubulin, GFAP, and β-actin for all APOE lines, with pooling of 5 organoids per independent differentiation batch. (**A**) APOE4/4: lanes 1, 3-8; APOE3/3: lanes 9-14; Molecular weight (MW) ladder: lanes 2, 15. Molecular weight indicated in KDa. Note that membranes were stripped once and reprobed for β-actin, allowing quantification of band intensity relative to their corresponding β-actin by computerized densitometry (Image Lab 6.1, Biorad), with APOE4/4 organoids represented in red and APO3/3 organoids in blue. Identity of each lane: 1, 4, 5: TOB0064; 3: TOB0121; 6-8: TOB0002c3; 9-11: MBE2960c1; 12-14: MBE2968c1. Each triplicate of APOE lines are from independent differentiation batches as follows: batch 1: 1, 6, 9, 12; batch 2: 4, 7, 10, 13; batch 3: 3, 5, 8, 11, 14. Expected sizes: PAX6 (46-48 kDa), β-tubulin (50-55 kDa), GFAP (37-50 kDa), β-actin (42 kDa). (**B**) APOE4/4: lanes 2-4; APOE3/3: lanes 5-9; molecular weight (MW) ladder: lanes 1, 10. Identity of each lane: 2-4: TOB0064; 5-7: iTOB0064_2_; 8-9: iTOB0064_3_. Note that membranes were stripped and reprobed as follows: first membrane PAX6, followed by β-actin; second membrane β-tubulin, followed by β-actin; third membrane GFAP followed by β-actin; fourth membrane APOE followed by β-actin. Independent differentiation batches as follows: batch 1: 2, 5, 8; batch 2: 3, 6, 9; batch 3: 4, 7. (**C-D**) Quantification of APOE protein by ELISA in organoid lysates (**C**) and conditioned media (**D**) in 6-month-old organoids. Lysates: MBE2960 n=7 (c1=3, c2=2, c3=2), MBE2968 n=6 (c1=2, c2=4), TOB0002 n=4 (c3), TOB0064 n=6, iTOB0064 n=5 (c1=3, c2=2). Supernatants: MBE2960 n=3 (c1), MBE2968 n=3 (c2), TOB0002 n=3 (c3)TOB0064 n=5, iTOB0064 n=5 (c1=3, c2=2). (**E**) Western blots of APOE for TOB0064 and iTOB0064_2_, with pooling of 5 organoids per independent differentiation batch. Membranes were stripped and reprobed with β-actin, allowing quantification of band intensity relative to their corresponding β-actin by computerized densitometry (Image Lab 6.1, Biorad). Molecular weight indicated in KDa. APOE4/4: lanes 2-4; APOE3/3: lanes 5-9; molecular weight ladder: lanes 1, 10. Identity of each lane: 2-4: TOB0064; 5-7: iTOB0064_2_; 8-9: iTOB0064_3_. Independent differentiation batches as follows: B=batch 1: 2, 5, 8; batch 2: 3, 6, 9; batch 3: 4, 7. Data are expressed as mean ± SEM. Statistical significance was evaluated using the one ANOVA followed by the Tukey’s test for multiple comparisons (**A**, **C**) or t-test (**B**, **D**, **E**) and established at *p<0.05, **p<0.01.

It has been shown that *APOE4/4* individuals produce less APOE than those with an *APO3/3* genotype [15]. We investigated whether the same could be observed in our cerebral organoids, since previous studies produced conflicting results [21, 25]. For this purpose, we used APOE3 and APOE4 iPSC lines to quantify APOE in 6-month-old cerebral organoids by ELISA (Figure 3C, D). As per above, these experiments were performed using independent lots of differentiation in order to take into account batch variations, with 5-6 organoids per batch. APOE was detected in lysates of all cell line-derived organoids, in similar amounts and with little variations between cell lines and between batches of differentiation, with the exception of the iTOB0064 samples which had a higher variability between batches (Figure 3C). In the unrelated iPSC lines, levels of APOE in organoid lysates were similar, regardless of the *APOE* genotype. The most amount of APOE was secreted, with a noticeable heterogeneity between unrelated cell lines and across batches of differentiation (Figure 3D). In the conditioned media, the APOE amounts were variable with potential differences between *APOE* genotypes (Figure 3D). However, given the variation in cellular content reported between independent cell lines (Figure 3A, B), it is possible that the levels of APOE recorded were correlated to an heterogeneity of cell type proportions rather than *APOE* genotypes. To accurately address this point, we used the isogenic lines TOB0064 and iTOB0064, as they showed similar cell type content, to assess secretion of APOE. This was measured by quantifying its levels in organoid lysates as well as their conditioned media. We did not observe significant changes in APOE amounts between the two genotypes (Figure 3C, D). Similar results were obtained by western blot of APOE in the cell lysates, with no statistically different levels of APOE between the isogenic lines (Figure 3E). This data thus suggests that the *APOE* status does not significantly modify levels of expression of APOE within the six-month-old cerebral organoids.

We then quantified Aβ levels in the six-month-old cerebral organoids by ELISA. Both cell lysates and their respective supernatants were analysed, and data were presented for organoids from individual lines, obtained in independent batches of differentiation (Figure 4). Aβ_1-42_ and Aβ_1-40_ were detected in organoid lysates and their supernatants of all cell lines, with large inter-cell line and intra-cell line variabilities (Figure 4). Levels of Aβ_1-42_ and Aβ_1-40_ were not statistically significantly different between unrelated iPSC-derived organoids of the two *APOE* genotypes, whilst statistically significant variations of both peptides were observed in conditioned media from unrelated iPSC-derived organoids (Figure 4A). The Aβ_1-42_/Aβ_1-40_ ratio did not show difference between genotypes and independent cell lines in lysates and supernatants (Figure 4). To assess if the significant differences in secreted Aβ peptides were cell line- or APOE-dependent, we quantified their amounts using the isogenic organoids. Within these, levels of Aβ_1-40_ and Aβ_1-42_ were similar in both lysates and conditioned media between the two genotypes (Figure 4), hence demonstrating that the APOE status does not influence Aβ levels in six-month-old cerebral organoids.

**Figure 4.**
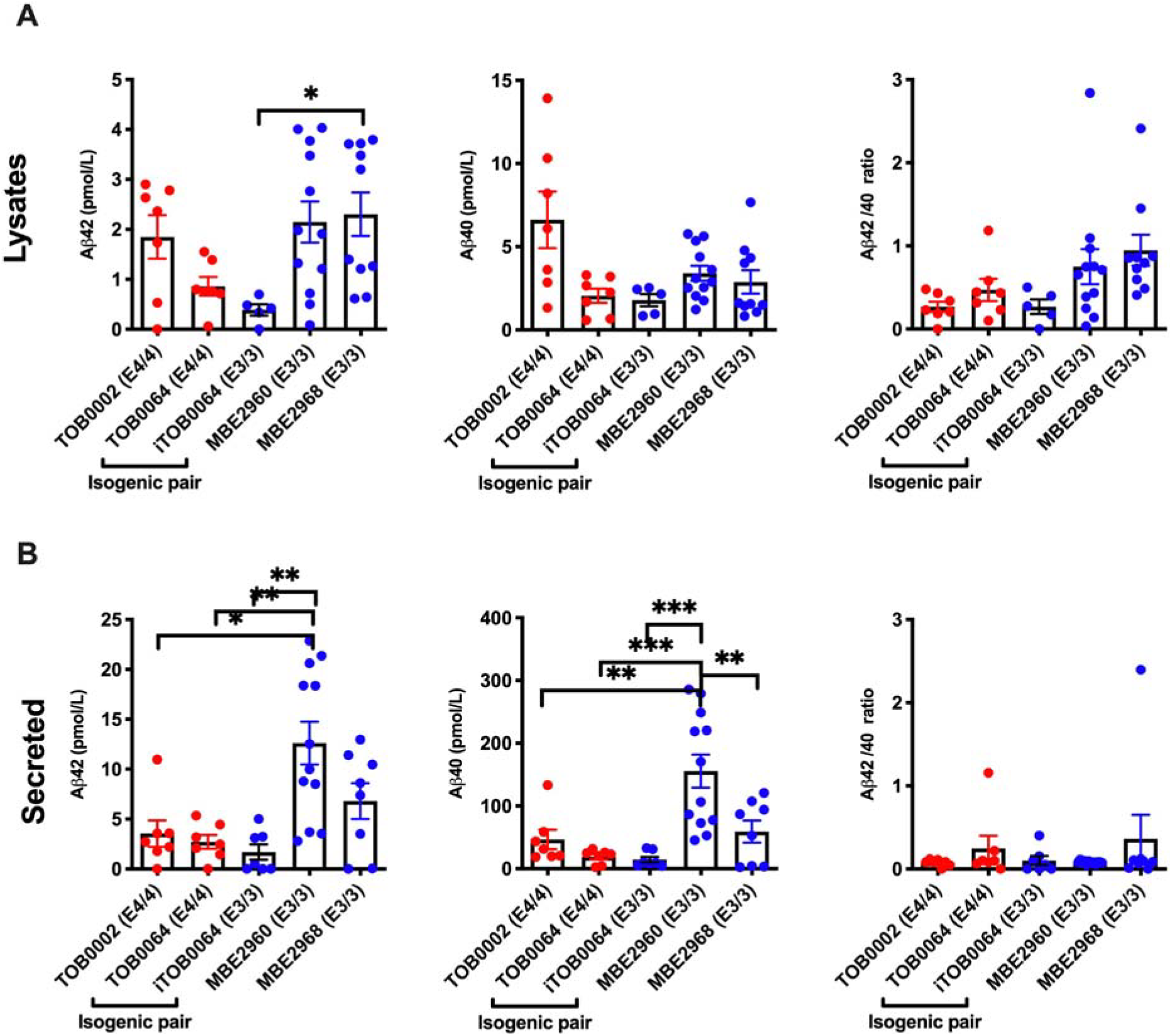
Quantification of Aβ production in cerebral organoids. ELISA of lysates (**A**) and supernatants (**B**) from six-month-old cerebral organoids, with APOE4/4 organoids represented in red and APO3/3 organoids in blue. (**A**) Lysates: TOB0002 n=7 (c3=5, c5=2), TOB0064 n=7, iTOB0064 n=5 (c2=3, c3=2), MBE2960 n=12 (c1=4, c2=4, c3=4), MBE2968 n=10 (c1=4, c2=6). (**B**) Secreted (supernatants): TOB0002 n=7 (c3=3, c5=3), TOB0064 n=7, iTOB0064 n=7 (c2=5, c3=3), MBE2960 n=12 (c1=3, c2=3), MBE2968 n=8 (c1=3, c2=3). Each dot represents the measurement of combined lysates or secreted media from 5-6 organoids from the same differentiation batch adjusted per 100 μg of total protein. Data are expressed as mean ± SEM. Statistical significance was evaluated using the one ANOVA followed by the Tukey’s test for multiple comparisons (for unrelated lines) or t-test (for comparison of TOB0064 and _i_TOB0064) and established at *p<0.05, **p<0.01, ***p<0.001.

Finally, we determined the phosphorylation status of Tau in the six-month-organoids by ELISA quantification of Tau, and its phosphorylated forms at S199, S396, and T231 in organoids’ lysates (Figure 5). These sites of phosphorylation were chosen as they correlate with different severities of cytopathology in AD [38]. Phosphorylation of Tau at S199 and S396 sites is associated with a late stage of neurofibrillary tangle development, and T231 phosphorylation with an early stage of neurofibrillary tangle development [38]. Tau was present in organoids from all cell lines, in similar levels and without statistically significant variation between the various lines and *APOE* genotypes (Figure 5A). However, as with previous assays, inter- and intra-cell line variabilities were observed. Tau phosphorylation levels were low, indicating that most of Tau was unphosphorylated in the organoids. As shown in Figure 5B-D, the ratio of phosphorylated Tau at different sites to Total Tau was consistently low, and heterogenous between cell lines and batches of differentiation. Some variations in pTau/Tau reached statistical significance between cell lines. However, given the overall variation of Tau levels between cell lines, it would be inaccurate to attribute these differences to specific genotypes. The inclusion of the isogenic CRISPR line provided clarity on these variations as both TOB0064 and iTOB0064 showed similar levels of total Tau (Figure 5A). In these organoids, the conversion of *APOE4* into *APOE3* was associated with a statistically significant decrease of the S199 phosphorylation levels (Figure 5B), and no modification of the phosphorylation at S396 and T231 (Figure 5 C, D).

**Figure 5.**
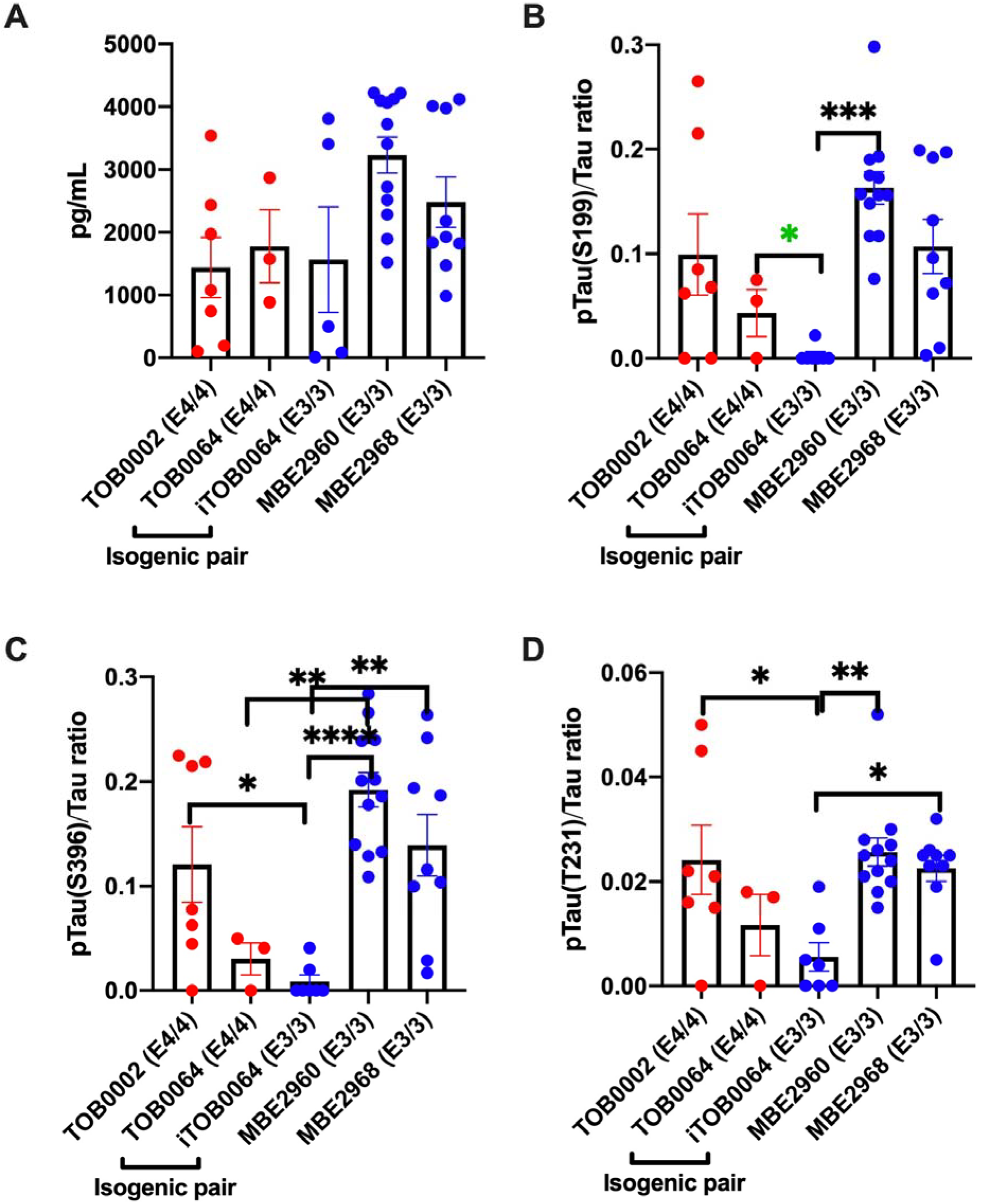
Quantification of Tau in cerebral organoids. ELISA of lysates from six-month-old cerebral organoids, with APOE4/4 organoids represented in red and APO3/3 organoids in blue. Lysates: MBE2960 n=12 (c1=4, c2=4, c3=4), MBE2968 n=9 (c1=4, c2=5),TOB0002 n=7 (c3=3, c5=2), TOB0064 n=3, iTOB0064 n=5 (c2=3, c3=2). Each dot represents the measurement of combined lysates from 5-6 organoids from the same differentiation batch. (**A**) Total Tau, (**B-D**) phosphorylated Tau at S199 (**B**), S396 (**C**) and T231 (**D**) sites adjusted to total Tau. Data are expressed as mean ± SEM. Statistical significance was evaluated using the one ANOVA followed by the Tukey’s test for multiple comparisons (for unrelated lines) or t-test (for comparison of TOB0064 and _i_TOB0064), and established at p<0.05, **p<0.01, ***p<0.001, ****p<0.0001. Green * indicates significance by t-test.

## Discussion

Potential advantages of using cerebral organoids over other approaches commonly used to model AD are that organoids show an organised structure, absent in neurospheres, and that organoids possess differentiated neurons resembling those of the brain regions implicated in AD, which is not the case with other constructs [3, 39, 40]. However, there are also important limitations to cerebral organoids that restrict their use as an *in vitro* model including AD. For instance, as cerebral organoids mimic development, they are composed of immature cells, and form a three-dimensional construct that is itself developing and maturing. Further, cells within organoids can follow an impaired cell type specification [41]. Additionally, because of their developing nature, organoids lack cell types normally observed in the fully formed brain, and have a different proportion of cell types than what would be observed in an adult brain. Moreover, given cerebral organoids are of ectodermal origin, they also lack the important microglial cells and vasculature normally present in a human brain. Hence, as AD is a condition for which symptoms present later in life, it is possible that these cannot be recapitulated at a cellular and molecular levels in an immature cell construct. Many reports have however clearly demonstrated the validity of using cerebral organoids to model specific aspects of AD in a dish [13]. Yet, it is important to ensure that phenotypes observed are truly linked to a given set of circumstances (such as a specific genetic risk factor) and not due to technical artefacts or limitations. Here, we sought to investigate if cerebral organoids from a small sample of unrelated individuals with either *APOE3/3* or *APOE4/4* genotypes could be robustly used to model key aspects of AD, namely APOE levels, Aβ production and Tau phosphorylation. We report the generation of *APOE3/3-* and *APOE4/4-* iPSC lines from unrelated individuals as well as a CRISPR-edited isogenic iPSC line with conversion of *APOE4/4* into *APOE3/3*. All lines were characterised and can now be added to the tools of reagents available to investigate the role of APOE in biological processes. Our data indicates that the APOE status does not significantly modify levels of expression of APOE within cerebral organoids at 6 months of culture. It is consistent with data reported in a recent study noting levels of APOE were not statistically different in 4, 8 and 12 weeks-old *APOE4/4* organoids compared to *APOE3/3* organoids [25]. Although this study concluded on increased levels of APOE withIn *APOE4/4* organoids when compared to *APOE3/3* organoids, the data presented did not show statistically significant variations in APOE levels between organoids of different APOE variants [25]. Although not considered a main source of APOE, neural progenitors can synthesize APOE [20]. Monocultures of neural progenitors derived from *APOE3/3*-iPSCs have been shown to produce higher levels of APOE than their isogenic *APOE4/4* cells [20], but those data were not confirmed in corresponding cerebral organoids [20], hence cannot be placed in context of our results. Interestingly, our data differs from those reported by [21] which demonstrated reduced levels of APOE in 6-month-old *APOE4/4* cerebral organoids when compared to their isogenic control *APOE3/3*. The apparent discrepancy of results could be due to intra-cell line variability as in their study, the authors did not assess cellular composition of cerebral organoids prior to measurement of APOE levels [21]. Nonetheless, LIn *et al* clearly demonstrated higher levels of APOE in monocultures of *APOE3/3*-iPSC-derived astrocytes when compared to *APOE4/4* cells [21]. It is thus possible that the relative amount of astrocytes within cerebral organoids, even after 6 months in culture, is not sufficient to adequately assess the specific influence of this cell type on the whole multicellular construct. It has already been demonstrated that cerebral organoids can be used to model Aβ and Tau pathologies in familial [42, 43] and sporadic (*APOE4/4*) forms of AD [21, 22, 25]. Our work clearly indicates that both Aβ secretion and Tau phosphorylation are present in cerebral organoids and that their presence in the organoids is not influenced by APOE. Indeed, although we observed Aβ secretion and low Tau phosphorylation, no statistical variation was observed between APOE3/3 and APOE4/4 cerebral organoids, with the exception of one site of phosphorylation of Tau which proved to be reduced when converting APOE4 into APOE3. This data is interesting as it differs from a recent study by [22] which described a role of APOE4 in Aβ production and in increasing the amount of phosphorylated Tau in human iPSC-derived neurons. Those variations in levels of phosphorylated Tau could be linked to the differences of cell cultures, in particular timing and cell types present. Indeed, cerebral organoids are composed of neural stem/progenitor cells, astrocytes and neurons, as opposed to homogenous cultures of neurons. Our work is however in accordance with [21] as no variation in Aβ secretion or Tau phosphorylation could be observed between APOE3/3 and APO4/4 organoids at 90 days. It also suggests that variations in those key pathological markers in neural tissue is likely due to their clearance by cells not present within the organoids, such as microglial cells, or by more mature cell types than those observed in the organoids. Given the presence of Aβ and Tau in amounts readily quantifiable by ELISA, cerebral organoids present a real potential to develop a new model in which to screen for modifiers of AD key pathological markers.

As we confirm the presence of Aβ and Tau in 6-month-old cerebral organoids with *APOE3/3* and *APOE4/4* genotypes, we also highlight the variability in these amounts which seems correlated to cell lines and batches of differentiation. This observation is consistent with the variability in cell type proportion between organoids from different lines and different batches of differentiation. It thus clearly highlights the importance of including isogenic lines to decrease variability in cell line behaviours but also the need of optimising protocols able to reduce the variability linked to experimental batches. Outside the scope of this study, but clearly of relevance, is the optimisation of automation of cell culture processes to streamline and improve consistency of methods.

## Acknowledgements and Funding

We thank Sophie Chevalier and Vikrant Singh for technical assistance. This work was supported by grants from the Yulgilbar Alzheimer’s Research Program, the DHB Foundation, the Brain Foundation, Dementia Australia, a National Health and Medical Research Council (NHMRC) Practitioner Fellowship (AWH), an NHMRC Senior Research Fellowship (AP, 1154389), the University of Melbourne and Operational Infrastructure Support from the Victorian Government.

## Author contributions

DH: concept and design, experimental work, interpretation of data, financial support, writing of manuscript, final approval of the manuscript; LR, MD, LG, HHL: experimental work, interpretation of data, final approval of the manuscript; ALC: provision of samples, final approval of the manuscript; AWH: provision of biopsies, interpretation of data, final approval of the manuscript; AP: concept and design, interpretation of data, financial support, writing of manuscript, final approval of the manuscript.

## Conflict of interest/disclaimers

None

## Supplementary Figures

**Figure S1.**
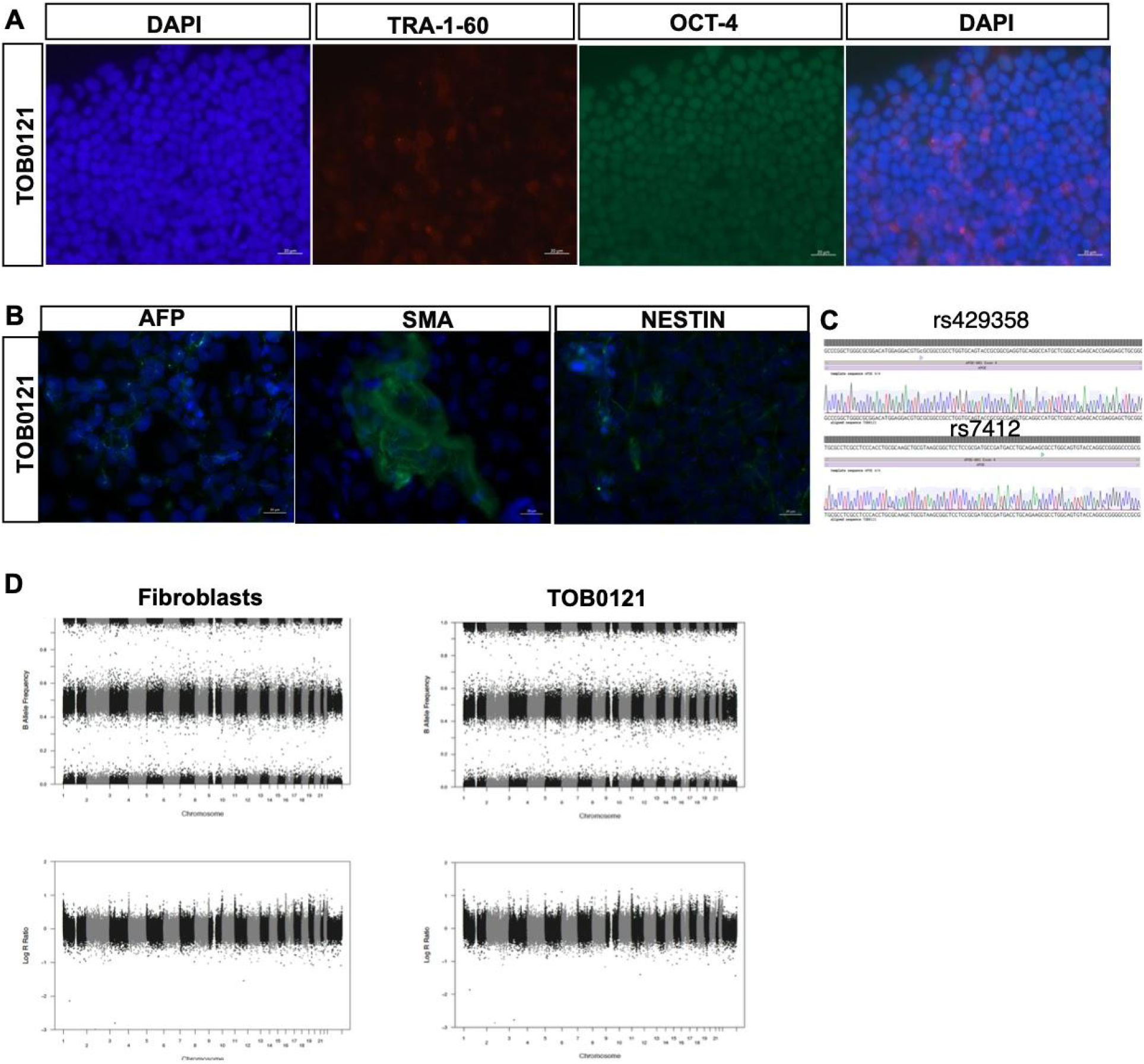
Characterisation of TOB0121. (**A**) **Generation of iPSC lines.** Representative images of TOB0121 showing expression of the pluripotency markers TRA-1-60 and OCT4, with DAPI counterstained and merged. Negative IgG controls are also displayed. (**B**) **Representative germ layer immunostaining of TOB0121.** TOB0121 demonstrates pluripotency by positive staining for markers of each embryonic germ layer; endoderm (AFP), mesoderm (SMA) and ectoderm (NESTIN). (**C**) ***APOE* genotyping.** Sanger sequencing of *rs429358* and *rs7412* in TOB0121 confirms *APOE4/4* status. (**D**) **Copy Number Variation Analysis of TOB0121 in their original fibroblasts and iPSCs (p8).** Each panel shows the B allele frequency (BAF) and the log R ratio (LRR). BAF at values others then 0, 0.5 or 1 indicate an abnormal copy number. Similarly, the LRR represents a logged ratio of “observed probe intensity to expected intensity”. A deviation from zero corresponds to a change in copy number.

**Figure S2.**
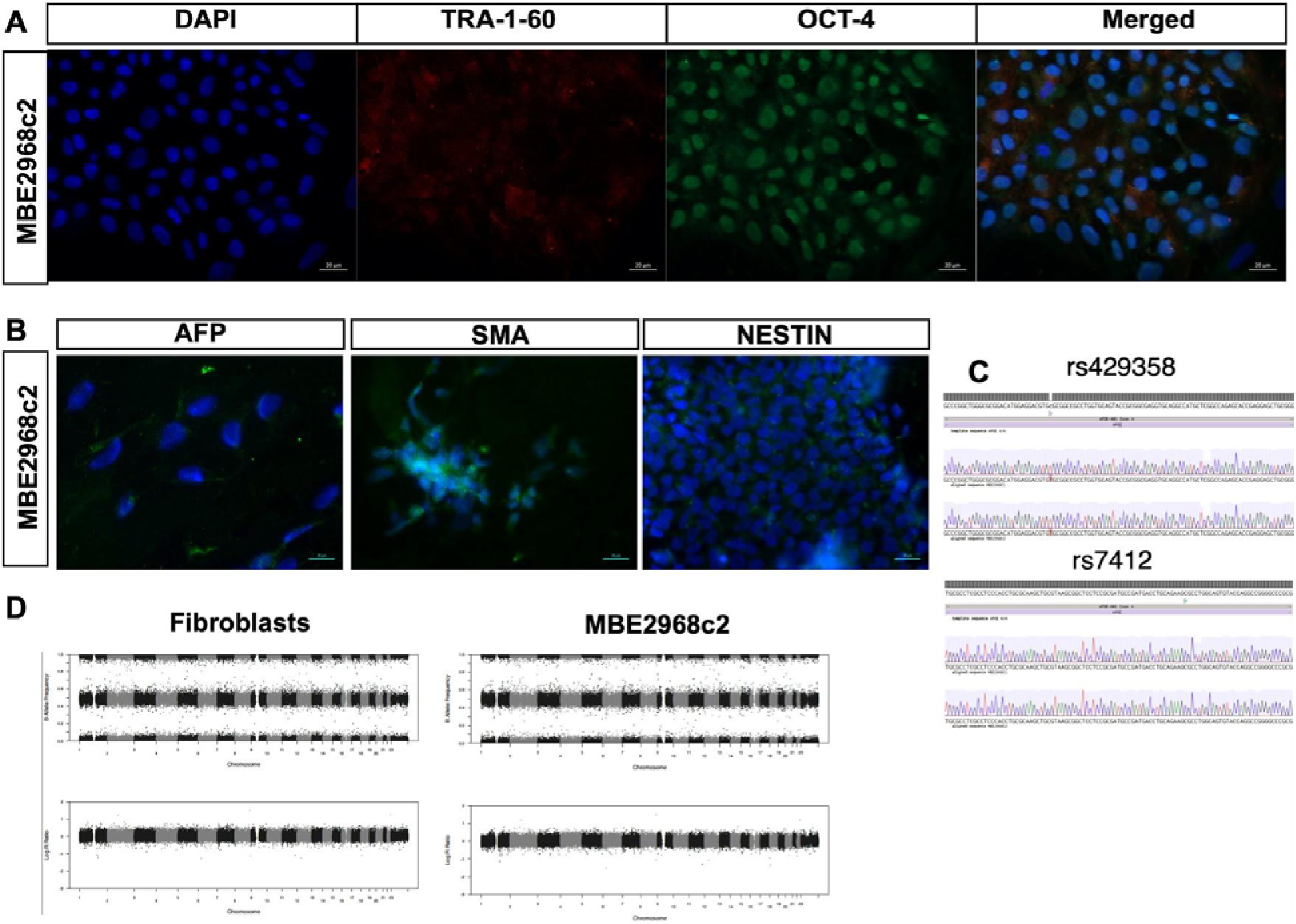
Characterisation of MBE2968. (**A**) **Generation of iPSC lines.** Representative images of TOB2968 clone (c) 2 showing expression of the pluripotency markers TRA-1-60 and OCT4, with DAPI counterstained and merged. (**B**) **Representative germ layer immunostaining of MBE2968.** MBE2968 c2 demonstrates pluripotency by positive staining for markers of each embryonic germ layer; endoderm (AFP), mesoderm (SMA) and ectoderm (NESTIN). (**C**) ***APOE* genotyping.** Sanger sequencing of *rs429358* and *rs7412* in MBE2968 c1 and c2 confirms *APOE3/3* status. Note that c1 genotype was already published in [23] (**D**) **Copy Number Variation Analysis of MBE2968 in their original fibroblasts and iPSCs (p8).** Each panel shows the B allele frequency (BAF) and the log R ratio (LRR). BAF at values others then 0, 0.5 or 1 indicate an abnormal copy number. Similarly, the LRR represents a logged ratio of “observed probe intensity to expected intensity”. A deviation from zero corresponds to a change in copy number.

**Figure S3.**
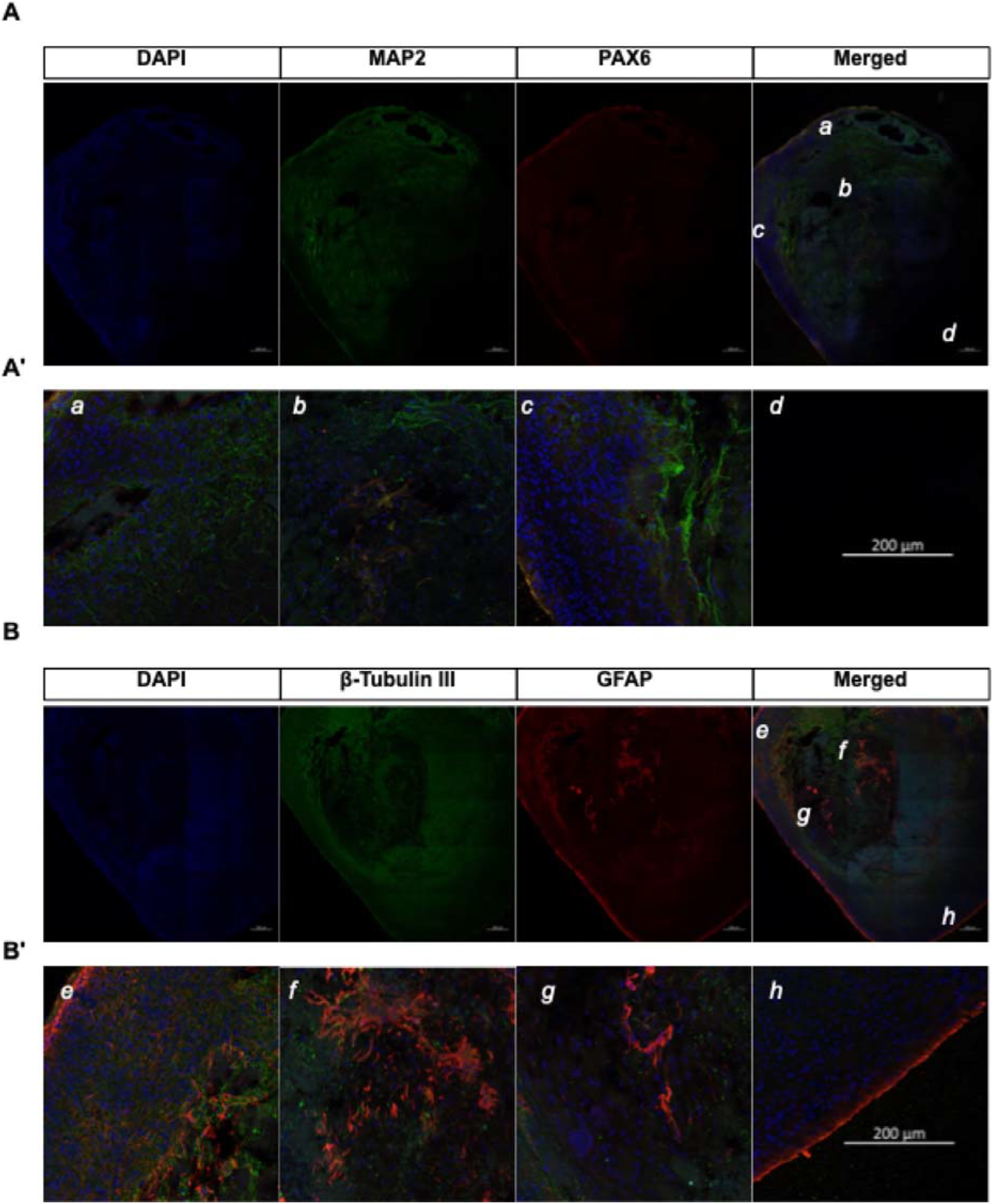
Representative immunostaining of 6-month-old cerebral organoids. Tile (**A, B**) and single (**A’**, **B’**) images of immunostaining of sectioned TOB0002 cerebral organoids for MAP2 with PAX6 (green and red respectively, **A, A’**), and β-Tubulin III with GFAP (green and red respectively, **B, B’**), counterstained with DAPI (blue) and merged (**A’, B’**). Images are representative of organoids obtained with all cell lines. Scale bars: 200 μm

**Figure S4.**
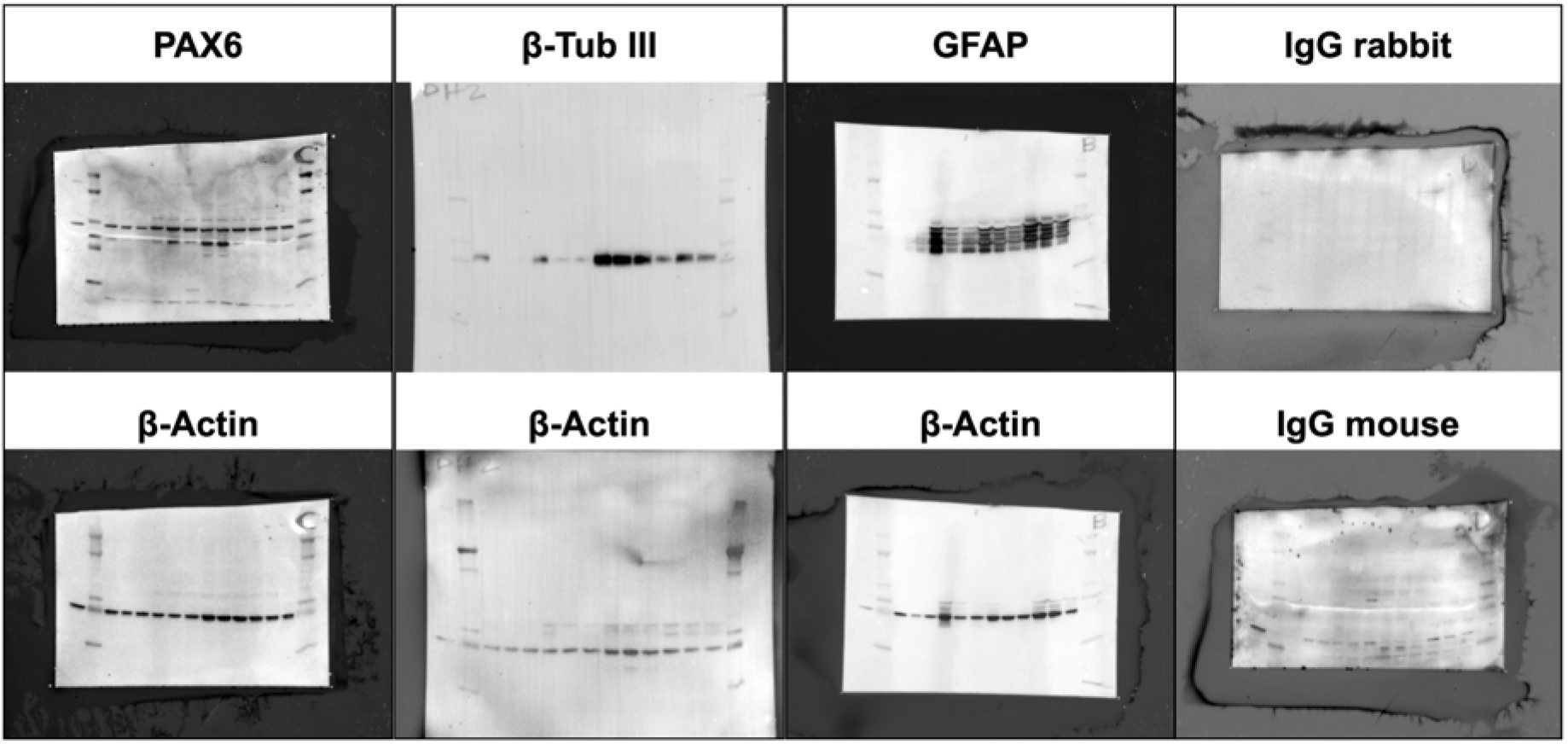
Western blots, uncropped membranes. Uncropped membranes for PAX6, GFAP, β-tubulin and β-actin negative isotype controls. Note that membranes were stripped and reprobed one time in the following order: PAX6 followed β-actin; β-tubulin followed by β-actin; GFAP followed by β-actin; rabbit IgG isotype control followed by mouse IgG isotype control. Relates to Figure 3A.

**Figure S5.**
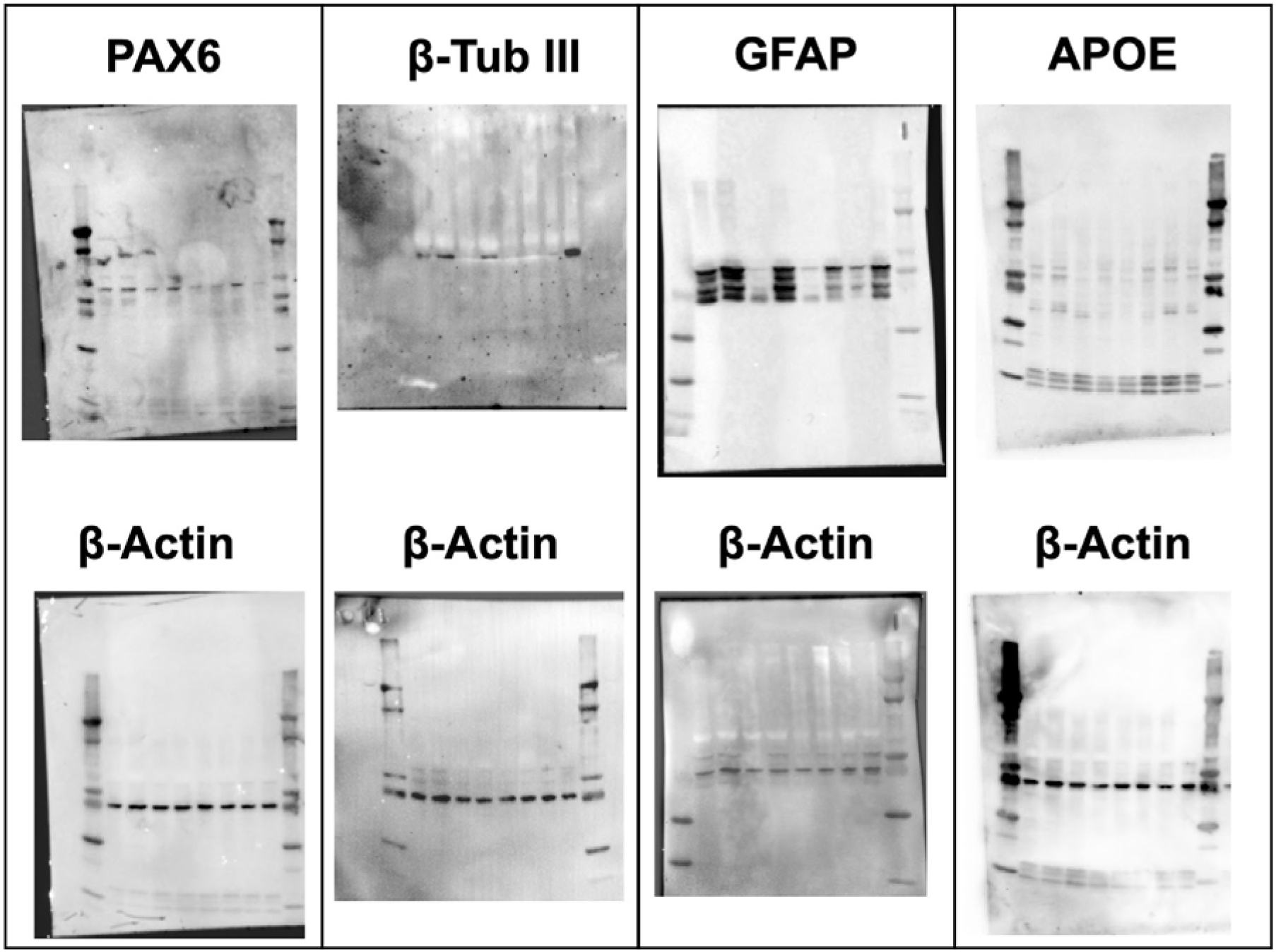
Western blots, uncropped membranes. Uncropped membranes for PAX6, GFAP, β-tubulin and β-actin. Note that membranes were stripped and reprobed one time in the following order: PAX6 followed by β-actin; β-tubulin followed by β-actin; GFAP followed by β-actin. Relates to Figure 3B, E.

